# Structural-mechanical remodelling of GDP-microtubules by kinesin

**DOI:** 10.1101/101022

**Authors:** Daniel R. Peet, Nigel J. Burroughs, Robert A. Cross

## Abstract

Kinesin-1 is a nanoscale molecular motor that walks towards the fast growing (plus) ends of microtubules (MTs), hauling molecular cargo to specific reaction sites in cells. Kinesin-driven transport is central to the self-organisation of eukaryotic cells and shows great promise as a tool for nano-engineering^1^^,^^2^. Recent work hints that kinesin may also play a role in modulating the stability of its MT track, both *in vitro*^3^^-^^5^ and *in vivo*^6^, but results are conflicting^7^^-^^9^ and mechanisms are unclear. Here we report a new dimension to the kinesin-MT interaction, whereby strong-state (ATP-bound and apo) kinesin-1 motor domains inhibit the shrinkage of GDP-MTs by up to 2 orders of magnitude and expand their lattice spacing by ~1.6%. Our data reveal an unexpected new mechanism by which the mechanochemical cycles of kinesin and tubulin interlock, allowing motile kinesins to influence the structure, stability and mechanics of their MT track.

---

During stepping, kinesin motor domains cycle through a series of nucleotide-specific conformations^10^^,^^11^. We tested different nucleotide states of kinesin-1 motor domains to quantify their influence on MT stability. We attached fluorescent MT ‘seeds’ to the inside of a flow chamber via biotin-NeutrAvidin linkages and flowed in GTP-tubulin, causing dynamic MTs to grow from the seeds (Fig. 1a). We then initiated MT depolymerisation by washing out tubulin, whilst simultaneously flowing in kinesin-1 motor domains. We used a kinesin motor domain mutant (T93N) as a stable homologue of the nucleotide-free (apo) state of the motor^12^, and compared this to wild-type (WT) motor domains (K340) in the presence of different nucleotides. Strong-state kinesin binds tightly and stereospecifically to MTs^13^. We found that T93N reduced MT shrinkage to 1% of the control rate (Fig. 1b,c) and that AMPPNP-WT kinesin had a similar effect, whilst weak-state (ADP-bound) kinesin had no detectable effect (Fig. 1c). We conclude that strong-state kinesin powerfully inhibits the shrinkage of GDP-MTs.

**Figure 1.**
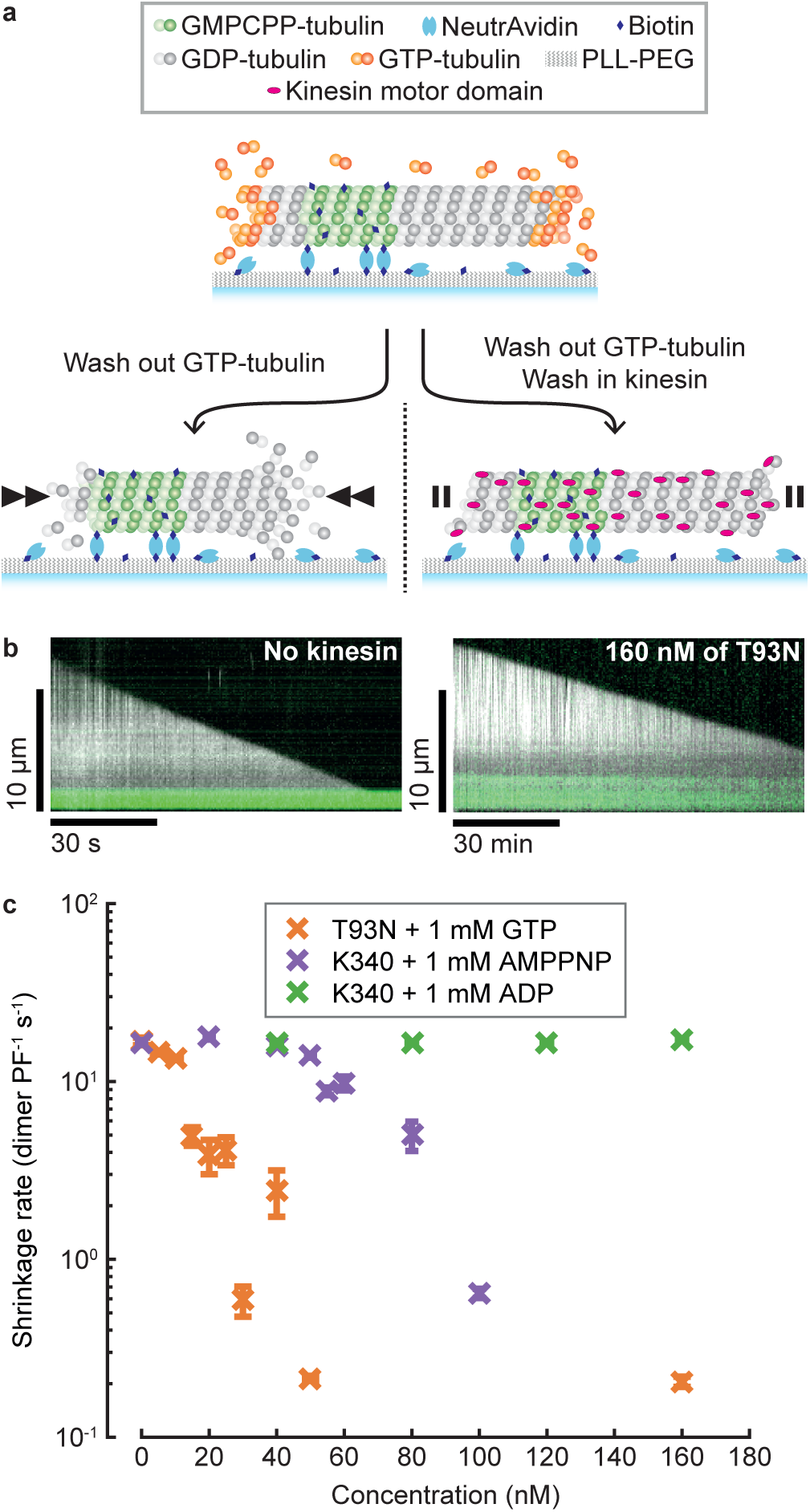
Strong-state kinesins inhibit GDP-MT shrinkage. **a,** Schematic representation of a tubulin depletion assay. Dynamic MTs shrink rapidly when GTP-tubulin is depleted (left) unless bound to strong-state kinesins (right). **b,** Representative kymographs of MTs shrinking in the absence (left) and presence (right) of T93N. Note the different time scales. Dynamic MTs are shown in white (dark-field) and fluorescent seeds in green (epi-fluorescence). **c,** Shrinkage rates of MTs bound to kinesins in distinct nucleotide states. GTP was included with T93N only. Error bars are mean ± SEM. 14 ≤ *n* ≤ 53 for all conditions.

Next, we bound MTs to a kinesin-coated coverslip in a flow chamber, triggered depolymerisation by washing out residual GTP-tubulin, and again observed MT shrinkage. Geometric constraints suggest that in this arrangement, at most 5 protofilaments can bind to the kinesin surface (Fig. 2a). Despite this, entire MTs were stabilised (Fig. 2c). We then flowed solutions through the channel in 2 steps (Fig. 2b). First, ADP was flowed in, reducing the fraction of kinesins in a strong state and increasing the MT shrinkage rate (Fig. 2c, Supplementary Movie 1). By titrating the ADP concentration, we found that MT shrinkage rates could be fine-tuned over 2 orders of magnitude (Fig. 2d, Supplementary Table 1). Comparing the inhibition of MT shrinkage by kinesin in solution (maximally 0.21 ± 0.02 (25) dimer PF^−1^ s^−1^ (mean ± SEM (*n*))) with that of the kinesin surface (0.06 ± 0.01 (6) dimer PF^−1^ s^−1^ in the presence of 400 nM ADP) shows that surface immobilisation enhances the stabilising effect of kinesin, despite its binding being restricted to only a subset of protofilaments.

**Figure 2.**
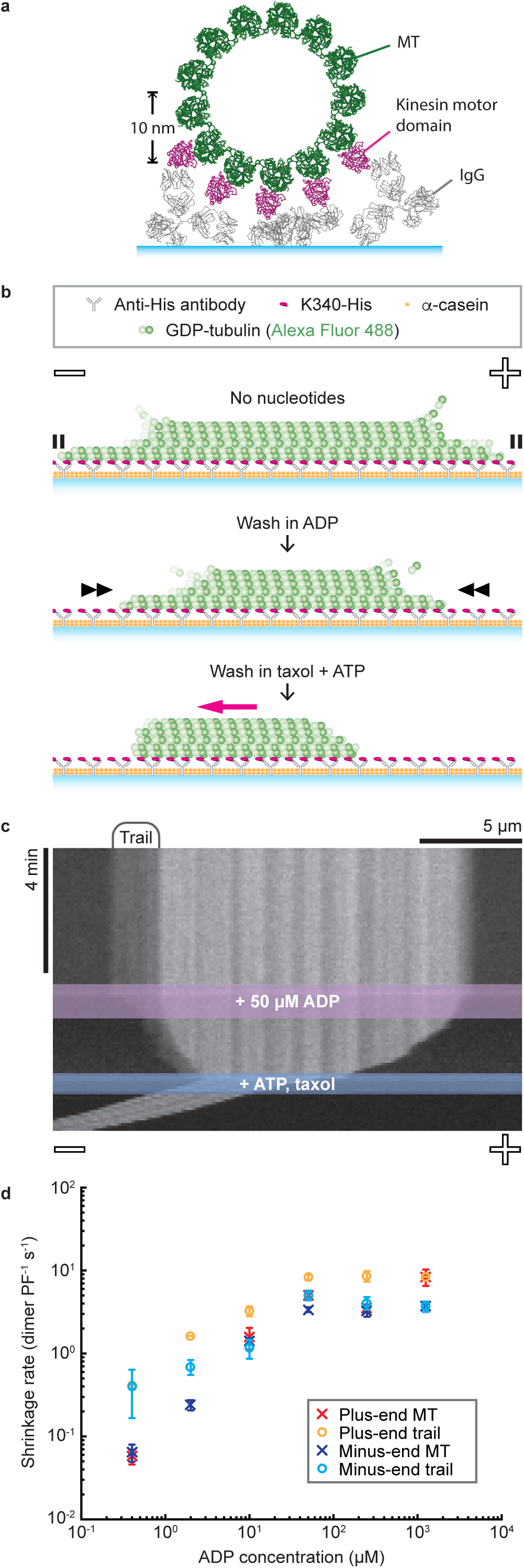
MTs are stabilised when kinesins bind to one side of the lattice. **a,** Cross-sectional view of a kinesin-clamp assay, showing IgG (PDB:1IGY) and kinesin-bound MT (PDB:4UXT) structures to provide scale. **b,** Schematic of a kinesin-clamp assay. **c,** Representative kymograph of a MT in a kinesin-clamp. **d,** Average shrinkage rates of MTs and their trails. Error bars are mean ± SEM, reflecting inter-MT variability. *n*-values are given in Supplementary Table 1.

Frequently, faint fluorescent trails were visible on the kinesin-coated surface in the wake of retreating MT tips. These shrank endwise upon addition of ADP, suggesting that their tubulin is still assembled into protofilaments (Fig. 2c, Supplementary Movie 1). Trails are tapered, and fluorescence intensity analysis (Fig. 3a, Supplementary Methods and Supplementary Fig. 1-2) indicates that at their tips they contain 2-3 protofilaments (Fig. 3b). On average, trails can shrink faster than their MT stem because they appear transiently, typically forming, lengthening and retracting multiple times during the shrinkage of each surface-attached MT (Supplementary Fig. 3). As a final step in these experiments, we flowed in a buffer containing taxol and ATP, triggering kinesin-driven sliding to reveal the MT polarity.

**Figure 3.**
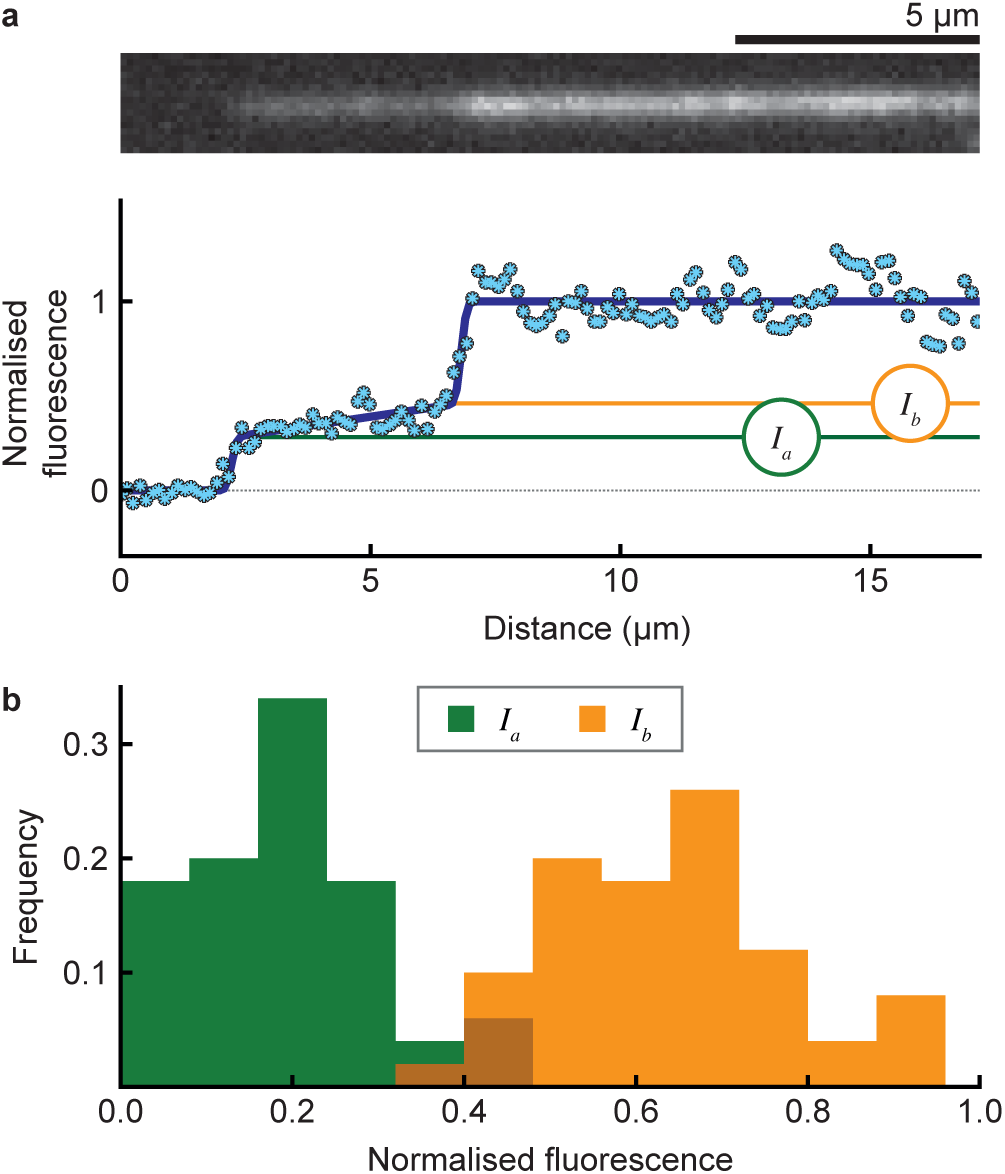
Subsets of protofilaments are stabilised by a kinesin-coated surface. **a,** Model fit to the intensity profile of a MT tip (bottom) with the associated fluorescence image (top). (I_a_) is the intensity at the tip and (I_b_) at the base of the trail. **b,** Histogram of (I_a_) and (I_b_) values for (*n* = 50) MTs, normalised to the intensities of their parent MTs in the no-nucleotide phase of the experiment. Mean ± SD is 0.19 ± 0.11 and 0.64 ± 0.13 for I_a_ and I_b_, respectively.

Why does a kinesin-coated surface stabilise MTs but also cause them to split? Taxol-stabilised MTs have recently been shown to split on a kinesin-coated surface^14^. However, ATP-driven kinesin motility was essential to this process, which is not the case for our GDP-MTs. Several strands of evidence suggest that kinesin binding can change the lattice conformation and mechanics of MTs. A kinesin-coated surface has been reported to reduce the Young’s modulus of taxol-stabilised MTs^15^. Structural changes have also been reported following kinesin binding to taxol-stabilised^16^ and GMPCPP-bound MTs^17^. Furthermore, the longitudinal compaction of the MT lattice that accompanies GTP hydrolysis is reduced in kinesin-bound MTs^18^, suggesting that kinesin influences the longitudinal spacing between tubulin subunits in the MT lattice. We therefore hypothesised that kinesin binding modifies the axial spacing between GDP-tubulin subunits in the MT lattice. Binding kinesins to one side of a MT, as in our surface assay, would then change the lattice spacing on that side but not the other, causing shear stress that could facilitate splitting.

To test the idea that kinesin binding stabilises a distinct conformation of the MT lattice, we used hydrodynamic flow to bend tethered dynamic MTs, thereby expanding the MT lattice on the convex side and compacting it on the concave side (Fig. 4, Supplementary Movie 2). We supplemented T93N into the flow and observed the mechanical response of MTs upon stopping the flow. In the absence of kinesins, the MTs quickly recoiled to a straight conformation and rapidly depolymerised. Remarkably, low concentrations of T93N (15-30 nM) blocked this recoil, effectively locking the GDP-MTs in a curved conformation as well as inhibiting their shrinkage. These data heavily suggest that indeed strong-state kinesins preferentially bind and stabilise a distinct longitudinal lattice spacing of GDP-MTs.

**Figure 4.**
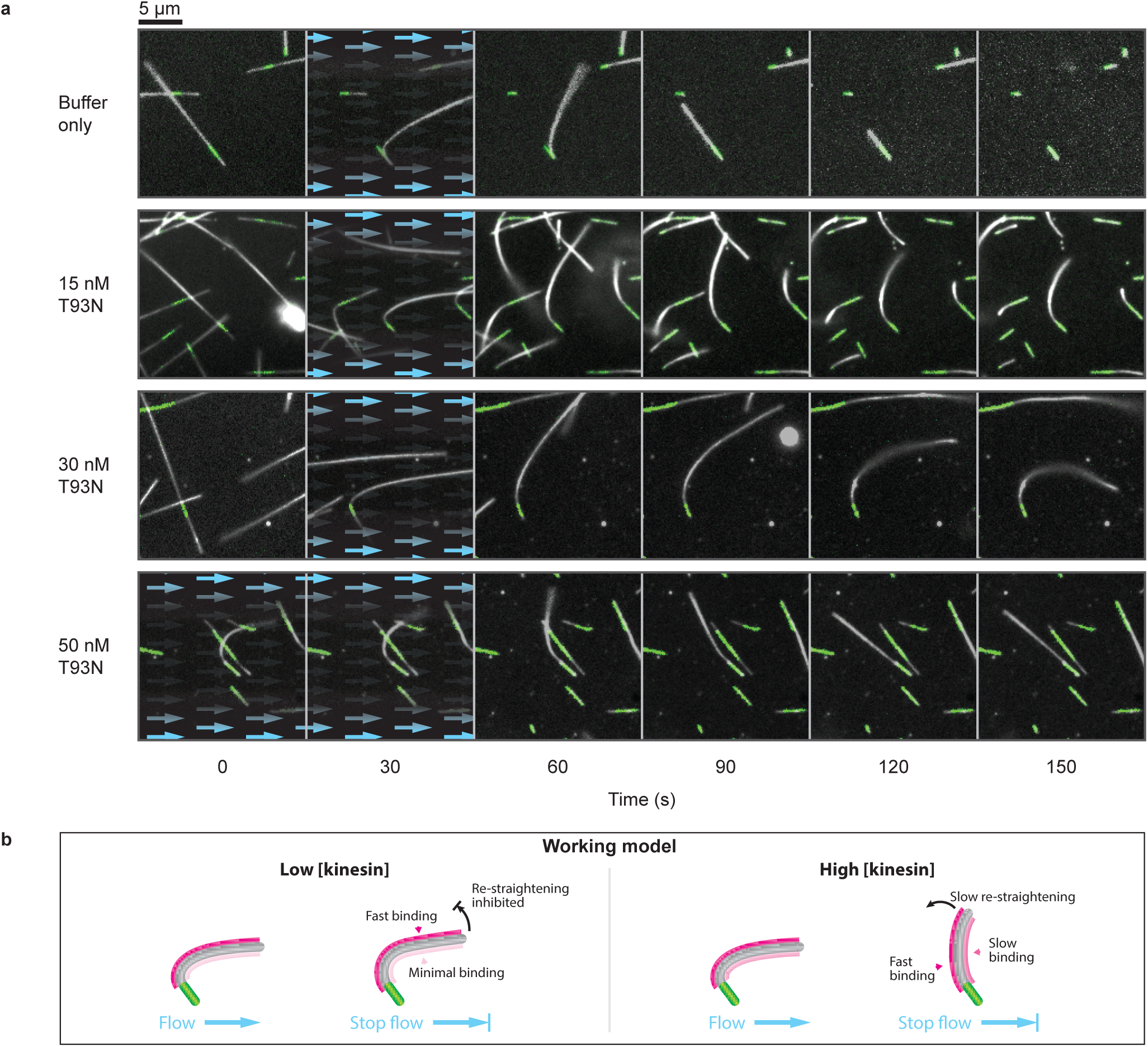
Nucleotide-free motor domains can bend-lock MTs. **a,** Time-lapse images of MT bending experiments for a range of kinesin concentrations. Blue arrows highlight the presence and direction of fluid flow. Dynamic MTs appear white (dark-field) and fluorescent seeds are marked in green (epi-fluorescence). Each condition was tested twice on independent occasions. MTs shown here have been selected for having similar orientations. A more extensive selection is given in Supplementary Movie 2, which shows a complete range of orientations and lengths. **b,** Working model. We propose that kinesin binds preferentially to the stretched (convex) side of the MT and also stabilises this expanded region of the MT lattice. At high kinesin concentrations, the convex side of the MT would quickly saturate. Binding would also occur slowly on the concave side, causing this side to expand, progressively restraightening the MT.

We noticed that in the presence of higher concentrations of kinesin (> 50 nM), the curved MTs tended slowly to re-straighten. To explain this observation, we speculate that strong-state kinesins bind preferentially *but not exclusively* to one side of curved MTs. At high kinesin concentrations, the favoured side of the MT would then quickly become fully occupied, while binding would continue more slowly on the unfavoured side, ultimately driving the MT back into a straight conformation (Fig. 4b). Kinesins have previously been reported to bind preferentially to GTP-MTs, which have a longer lattice spacing than GDP-MTs^19^. In light of this, we postulated that the binding of strong-state kinesins drives an increase in the lattice spacing of GDP-MTs.

In order to directly test this model, we grew dynamic MTs from surface-tethered fluorescent seeds as previously but this time we capped their exposed tips with fluorescent GMPCPP-tubulin (Fig. 5a). We aligned the MTs using pressure-driven microfluidics. The GDP-MTs and their stabilised caps were fluorescently labelled in different colours and imaged using TIRF microscopy. The microfluidics system allowed us to maintain a constant flow rate whilst introducing either 200 nM of WT kinesin motor domains or 1 mM ADP into the flow. When apo-kinesin was introduced, the GDP-bound segment of the MT lengthened as predicted (Fig. 5b, Supplementary Movie 3).

**Figure 5.**
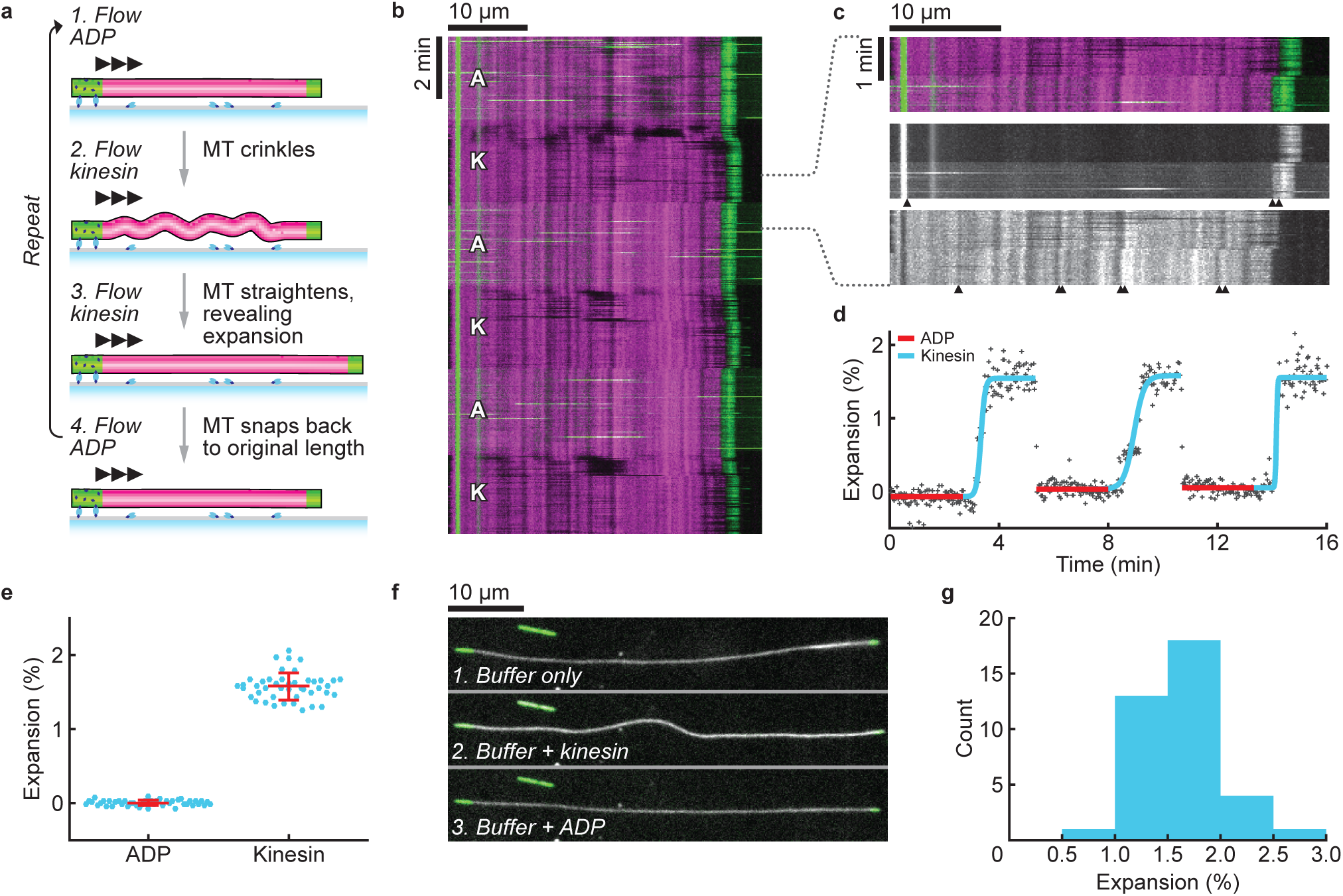
Kinesin increases the lattice spacing of GDP-MTs. **a,** Fluorescently labelled GDP-MTs (magenta) were grown from surface-bound GMPCPP-tubulin seeds (green) and then capped with GMPCPP-tubulin. Buffer was flowed through the channel at a constant rate and while switching between ADP and kinesin containing solutions. MTs transiently crinkled when kinesin was added, before straightening to reveal they had lengthened. Addition of ADP caused immediate recoil of the MTs to their original lengths. **b,** Representative kymograph of a MT changing length as the flow switched between ADP (A) and kinesin (K) containing buffers. A constant volumetric flow rate was maintained throughout the experiment (74.3 ± 0.3 μl/min; mean ± SD, 480 time points). This translated to a flow velocity of 248 ± 43 μm/s (mean ± SD, *n*=43) near to the surface, measured by tracking fluorescent beads that were included in the ADP buffer. **c,** Magnified region from panel **b** highlighting the contraction observed when switching from kinesin-containing to ADP-containing solutions. *Top*: merge. *Middle*: Green imaging channel (GMPCPP-tubulin and beads). Buffer exchange is visible due to the high background provided by the fluorescent beads (after ~50 sec). Arrowheads highlight the graduated movement along the MT during the exchange. The untethered tip retracts in <4 s (2 frames) upon switching to ADP. *Bottom*: Magenta imaging channel (GDP-tubulin). Fiducial markings reveal the expansion along the length of the MT. **d,** Expansion measured for the MT shown in panel **b**, obtained by scaling the fluorescence intensity profile of each time point (in the magenta imaging channel) to match the average profile when ADP is present. **e,** MT lattice expansion, showing 44 measurements from 16 MTs. Values are 0.00 ± 0.04 for ADP (control) and 1.58 ± 0.18 for kinesin (mean ± SD). **f,** Sequential images of a surface-stitched GDP-MT (dark-field image shown in white). Flow in this experiment was intermittent and images shown were taken in the absence of flow. When kinesin was added, the MT bowed to accommodate expansion of the MT lattice between the surface-bound points. The MT straightened and shortened upon addition of ADP. **g,** Expansion of surface-stitched GDP-MTs, given by the ratio of GDP-MT contour lengths with and without kinesin. Mean ± SD is 1.64 ± 0.39 (*n*=37).

TIRF microscopy visualises an optical section ~100 nm deep and MTs remained visible throughout the experiment, indicating that the flow constrained them within this 100 nm thick section. Strikingly, the MTs briefly crinkled upon switching from ADP to kinesin, causing them to dip in and out of the TIRF illumination (Supplementary Movie 4). This transient crinkling can also be seen in our initial MT bending experiments (Supplementary Movie 2). The crinkles progressively straightened (within 10-30 s) to reveal that the MT had expanded. Switching to ADP caused MTs to quickly recoil to their original length (<4 sec, 2 frames in our data) as the kinesin unbound (Fig. 5c, d). Quantification reveals that apo-kinesin binding to GDP-MTs increases their length by 1.6%, via a uniform axial expansion of the MT lattice (Supplementary Animation 1). The kinesin-induced lattice expansion appears to be fully reversible and the expand-and-recoil cycle can be repeated multiple times (Fig. 5d).

We additionally performed experiments where we used methylcellulose rather than flow to encourage the MTs to remain in focus, introducing a flow only intermittently to exchange buffers. Under these conditions MTs tended to become ‘stitched’ to the coverslip at sparse interaction sites. Introducing kinesin then caused the MTs to bow locally between these sites (Fig. 5f, Supplementary Movie 5), emphasising the expansion of the lattice. By measuring the change in contour length between the two stabilised MT caps, we confirmed that this bowing accommodates a 1.6% expansion of the GDP-MT lattice (Fig. 5g). MTs became more densely ‘stitched’ to the surface if multiple cycles were performed but this did not influence the measured expansion (Supplementary Fig. 4). When ADP was flowed through the channel, the MTs again recoiled to their original lengths (Fig. 5f).

Our data show that strong-state kinesin stabilises the GDP-lattice of dynamic MTs, and concomitantly increases their axial lattice spacing by 1.6%. Kinesins are known to bind to the intra-dimer interface of αβ-tubulin, away from the inter-dimer contacts of the MT lattice^10^^,^^11^^,^^17^^,^^18^^,^^20^.This suggests to us that kinesin binding allosterically modifies the conformation of GDP-tubulin, giving it properties more similar to GTP-tubulin. A 1.6% expansion equates to approximately 1.3 Å per 80 Å tubulin dimer, similar to the 1.7 Å difference observed by cryo-EM between GMPCPP-MT-kinesin and GDP-MT-kinesin structures^20^. Strong-state kinesins have previously been reported to alter the structure of both taxol-GDP-MTs^16^ and GMPCPP-MTs^17^. Moreover, a long-range, ATP-dependent cooperative effect has been described whereby the first few kinesins that bind facilitate subsequent binding events in the same region of the MT, again suggestive of a kinesin-induced conformational change^21^.

We envisage that the ability of strong-state kinesin to stabilise GDP-MTs by inducing a conformational change in their tubulin subunits provides at least a partial mechanistic explanation for the surface-bound depolymerisation trails and the bend-locking phenomenon reported here. Thus, a MT landing on and binding to a kinesin-coated surface would likely become stretched on its surface-bound side. This stretching would create shear stress in the lattice and potentially contribute towards formation of the trails observed in our kinesin-clamp experiments. Similarly, for the MT bend-locking, expanding the longitudinal MT lattice spacing by 1.6% exclusively on one side of the MT would be more than sufficient to account for the observed kinesin-stabilisation of curvature. Indeed, full occupancy on one side with zero occupancy on the other would produce a radius of curvature of 1.6 μm, far tighter than we observe in any of our post-flow data.

We have worked with kinesin-1, the best-studied kinesin, but it is possible that the mechanism we report here is common to other kinesins. Kif14 is a slow kinesin that binds to MTs in a rigor-like conformation and inhibits their shrinkage^22^. Kinesin-5 is reported to stabilise protofilament assemblies during MT growth^23^, potentially due to its strong state stabilising the polymer. Kip2^24^ and Kip3^25^ are also reported to dwell at MT ends and influence MT stability.

Our work reveals a specific action of strong-state kinesins in stabilising the GDP-lattice of dynamic MTs. MT activated ADP release creates a strong (apo) state and this process is affected by the tubulin and kinesin species^26^, by post-translational modifications^27^ and by the nucleotide state^28^ of the MT. Importantly, the residency of kinesin in the strong states is also influenced by mechanical force^29^. Such forces will arise *in vivo* wherever kinesins do mechanical work, for example at kinetochores, in MT bundles^30^, at cortical attachment sites^31^, and during the transport of vesicles against a resistance^32^. It will be important now to understand the role of these various effects in determining how kinesin motility may feed back on MT dynamics.

In conclusion, our data reveal a novel mechanism that allows kinesin-1 to feed back on the structure and stability of its MT track. Recent advances in the remote control of kinesin motility, such as photo-switchable fuels^33^, suggest the potential for precise spatial control of these effects.

## Methods

### Proteins and biochemical reagents

Tubulin was purified from pig brains as previously^3^ with additional steps as follows. Tubulin was polymerised in 50 mM PIPES, 1.2 mM MgSO_4_, 1 mM EGTA, 1mM GTP, 1 mM dithiothreitol (DTT) and 186 mg ml^−1^ glutamic acid and incubated for 60 min at 37 °C MTs were centrifuged in a TLA 100.3 rotor at 85,000 rpm for 20 min at 35 °C resuspended in K-PEM with 1 mM GTP, 1 mM MgSO_4_ and 1 mM DTT, cooled to 4 °C and centrifuged in a TLA 100.3 rotor at 85,000 rpm for 20 min. The supernatant was run through a Hiprep 26/10 desalting column into K-PEM buffer (100 mM PIPES, 1 mM MgSO_4_, 2 mM EGTA (Fisher); adjusted to pH 6.9 with KOH) and 20 μM GTP. Tubulin concentrations were determined using *E*_280_ = 105,838 M^−1^ cm^−1^.

X-rhodamine labelled tubulin was purchased from Cytoskeleton Inc. (USA). Alexa Fluor 488 (Molecular Probes) labelled tubulin was prepared using standard protocols^34^.

Kinesin was purified as previously^35^. Kinesin concentrations were determined using *E*_280_ = 15,300 M^−1^ cm^−1^.

Nucleotides were from Jena Biosciences (Germany). Other reagents were from Sigma (UK) unless stated otherwise.

### Bead-mPEG crosslinking

0.5 μm yellow-green carboxylated FluoSpheres (Thermofisher) were diluted to 1% solids and activated using 10 mg/ml 1-ethyl-3-(3-dimethylaminopropyl)carbodiimide (EDAC) in pH 6 MES buffer and mixed gently at room temperature for 30 min. Beads were then centrifuged, resuspended in 10 mg/ml of methoxypolyethylene glycol amine 750 (mPEG-amine) in pH 7.4 PBS and mixed gently at room temperature for 2 h. Adding 90 mM glycine quenched the reaction. After 30 min the beads were centrifuged and resuspended in 0.1% Tween20 in K-PEM 5 times and stored at 4 °C.

### Flow chamber assembly (for manual flow-through)

Flow chambers were assembled from 22x22 mm no. 1.5 glass coverslips (Menzel, Germany) and 76x26 mm 1-1.2 mm thickness glass slides (Menzel, Germany). Double-sided Scotch tape was sandwiched between the glass surfaces to form a 2 mm wide channel. The periphery of the chamber was further secured using nail polish excluding the channel ends, which were left open. Solutions were drawn through the channel by using grade 1 Whatman filter paper.

### Microfluidics

Microfluidic flow chambers were assembled by stacking the following: a 50x22mm no. 1.5 glass coverslip (Menzel, Germany), cleaned using the same protocol as the tubulin depletion assays; 81 μm thick double-sided adhesive tape (ArCare 90445, kindly provided by Adhesives Research), with a Y-shaped channel (two 8x0.75 mm inlets joining a 22x1.5 mm channel) cut out using a Silhouette Portrait plotter cutter; a 40x22mm cyclo-olefin polymer (COP) coverslip (ZF14-188, Zeon) with portholes cut out using a plotter cutter. Ports were assembled from ring magnets (8.16 mm OD x 3.5 mm ID, First4magnets) pressed into a 3D-printed ABS magnet holder that aligns the magnets with the portholes in the COP coverslip, which was then cast in polydimethylsiloxane (PDMS; Sylgard 184, Dow Corning) to fill the core of the ring magnets and form a 0.8 mm thick cushion to form the interface between the magnets at the COP coverslip. A 1.25 mm biopsy punch was used to bore holes through the PDMS-filled core of the magnets. 127 μm ID x 1.59 mm OD polyether ether ketone (PEEK) tubing was then inserted into the holes, forming a tight seal. The flow chamber was placed on a custom made ferromagnetic stainless steel (grade 430) microscope stage, which sealed the interface with the tubing by attracting the magnets and held the sample in position. Flow was driven using an MFCS-EZ, inlets were closed and opened using an L-switch, and the flow was monitored on the outlet using a size M flow unit (Fluigent). A function was written in Matlab (Mathworks) to provide fully integrated control of the microfluidics with the microscope.

### Tubulin depletion and MT bending assay

Coverslips were sonicated (600 W bath, Ultrawave) in 3% Neutracon detergent (Decon Laboratories, UK) for 30 min at 60 °C before undergoing extensive wash-sonication cycles in ultrapure water (18.2 MΩ resistivity). A flow chamber was then assembled, filled with 0.2 mg ml^−1^ PLL-PEG-biotin (SuSoS, Switzerland) and incubated for 30 min. The chamber was washed with K-PEM before adding 1 mg ml^−1^ NeutrAvidin (Thermo Fisher Scientific) for 5 min and washing again. MT seeds (polymerised by incubating 26 μM of 15% labelled Alexa488-tubulin and 1 mM GMPCPP in K-PEM at 37 °C for 25 min) were pelleted, diluted to ~ 60 nM, injected into the chamber and incubated for 5 min. After washing the chamber with K-PEM, dynamic MT extensions were grown from the seeds by flowing through with 15 μM tubulin, 1 mM GTP, GOC oxygen scavenger (4.5 mg ml^−1^ glucose, 0.2 mg ml^−1^ glucose oxidase, 35 μg ml^−1^ catalase, 0.5% (v/v) β-mercaptoethanol), 1 mg ml^−1^, 1 mg ml^−1^ BSA and 0.1% (v/v) Tween20 in K-PEM. MTs were allowed to grow for > 15 min at 25 °C prior to imaging with epifluorescence and dark-field illumination. Tubulin was then depleted by flowing through pre-warmed (25 °C K-PEM, supplemented with kinesin motor domains and nucleotides as described in the main text. MT bending was achieved by rapidly drawing solutions through the channel using Whatman filter paper.

### Kinesin-clamp assay

Fluorescence controls (colour-segmented stabilised MTs) were prepared by incubating 5 μM 30% labelled Alexa Fluor 288 tubulin and 0.2 mM Guanosine-5’-[(α,β)-methyleno]triphosphate (GMPCPP) in K-PEM at 37 °C for 60 min and pelleted in an airfuge (Beckman Coulter) at 25 psi for 10 min. The supernatant was discarded and the pellet resuspended in pre-warmed 5 μM 30% labelled X-rhodamine tubulin and 0.2 mM GMPCPP in K-PEM. MTs were left to anneal at room temperature then diluted 50-fold before use.

Coverslips were sonicated (600 W bath) at room temperature in a 1:1 solution of methanol and HCl for 30 min, then sonicated for 4 × 5 min in ultrapure water. Coverslips were then sonicated in 0.2 M KOH for 60 min, then sonicated for 5 × 5 min in ultrapure water. The coverslips were then spun dry using a Spin Clean (Technical video), incubated at 100 °C for 30 min and plasma-cleaned (PLASMA clean 4, ILMVAC) for 5 min. Coverslips were then silanised by immersion in 0.05% dimethyldichlorosilane in trichloroethylene for 60 min, washed in methanol, sonicated for 5 × 5 min in methanol and spun dry.

A flow chamber was assembled using a silanised coverslip and filled with 0.1 mg ml^−1^ anti-6xHistidine antibodies (372900) for 10 min. The surface was then blocked by filling the chamber with 0.5 mg ml^−1^ α-casein and incubating for 5 min. 75 nM K340 was then introduced and incubated for 10 min, after which the chamber was washed with 10 chamber volumes of K-PEM. Stabilised segmented MTs were then introduced. Unbound MTs were washed out immediately with K-PEM. Dynamic MTs, polymerised by incubating 50 μM 30% labelled Alexa Fluor 488 tubulin (same stock as used for fluorescence controls) and 1 mM GTP in K-PEM for 45 min at 37 °C were diluted 20-fold in warm (37 °C K-PEM and immediately flowed through the chamber. Next, 10 chamber volumes of warm K-PEM were flowed rapidly through the chamber. The sample was then imaged using epifluorescence on the dark-field microscope and ADP in K-PEM was introduced at the desired concentration. Once MTs had shortened sufficiently, 10 μM taxol and 200 μM ATP in K-PEM was flowed into the chamber.

### MT expansion assay with microfluidics

0.2 mg ml^−1^ PLL-PEG-biotin was pipetted into a microfluidic flow chamber and incubated for > 15 min before connecting to the pump and washing through each inlet with 50 μl of K-PEM. Dynamic MTs were then polymerised following the same initial steps as the tubulin depletion assay, except 20 μM of 10% x-rhodamine-labelled tubulin was used to grow the dynamic extensions, which were left for 1 h to polymerise, and buffer exchanges were performed using the microfluidic pump. MTs were then capped by incubating with 6 μM of 15 % Alexa-488 tubulin + 1 mM GMPCPP for 20 min before switching to 0.1% Tween20 + 2 mg/ml BSA for 5 min. The two pressure outlets of the pump were calibrated to achieve the same user-defined flow rate immediately before imaging with TIRF microscopy. K-PEM with 1X GOC, 0.1% Tween20, 0.02% methylcellulose (1500 cP) and 2 mg/ml BSA was then flowed through with 1 mM ADP + 0.01% mPEG-beads or 200 nM K340, switching between the two every 80 frames, cycling 3 times. Frames were acquired every 2 s. Beads were additionally flowed through immediately after the experiment ended and frames were acquired every 0.2 s for measuring the flow velocity (tracked manually). Images shown had the background subtracted in the green imaging channel by subtracting the median and applying a rolling ball (*r*=200) to each frame using Fiji/Imagej.

### Surface-stitched MT expansion assay

Coverslips were incubated in 1 M HCl at 50 °C for 12-15 hours, rinsed with ultrapure water twice, sonicated in ultrapure water for 30 min, sonicated in ethanol for 30 min, rinsed in ethanol and dried by spinning or spraying with nitrogen gas. MTs were then polymerised as in the MT expansion assay with microfluidics, except the MT seeds and caps were labelled to 30% with Alexa488 and the dynamic extensions were grown using unlabelled tubulin. MTs were capped for 10 min before washing the chamber with 100 μl of 0.1% Tween in K-PEM. Hereafter, each buffer contained GOC, 0.1% Tween20 and 0.02% methylcellulose (4000 cP) in K-PEM. The chamber was washed with 30 μl of buffer before imaging. During imaging, 40 μl of 200 nM K340 was flowed through manually, followed later by 40 μl of 1 mM ADP. Flowing through with 100 μl of buffer depleted the ADP, after which a new field of view could be imaged using epifluorescence and dark-field illumination and the kinesin and ADP flows repeated. We imaged no more than five times in a single flow channel.

### Dark-field/epifluorescence microscopy

Images were captured by an EM-CCD camera (iXon^EM^+DU-897E, Andor) fitted to a Nikon E800 microscope with a Plan Fluor 100x NA 0.5-1.3 variable iris objective. A custom-built enclosure with an air heater (Air-Therm ATX, World Precision Instruments) was used to keep samples at 25 °C. Dark-field illumination was achieved using a 100 W mercury lamp connected to the microscope via a fibre optic light scrambler (Technical video), cold mirror, 500-568 nm band-pass filter (Nikon) and a dark-field condenser (Nikon). A stabilised mercury lamp (X-cite exacte, Lumen Dynamics) provided illumination for epifluorescence, connected to the microscope with a light pipe. Motorised filter wheels (Ludl Electronic Products) housed the fluorescence excitation and emission filters: 485/20 and 536/40 for Alexa-488 and 586/20 and 628/32 for X-rhodamine (Chroma). Combined dark-field and fluorescence imaging was achieved using an FF505/606-Di01-25x36 dichroic mirror (Semrock) and electronic shutters to switch between illumination modes. The shutters, filter wheels and camera were controlled using Metamorph software (Molecular Devices).

### TIRF microscopy

Images were captured by an EM-CCD camera (iXon_3_ 888, Andor) fitted to a Warwick Open-Source Microscope (WOSM; wosmic.org), which was equipped with 473 nm (Cobolt) and 561 nm (Obis) laser lines and a Nikon 100x NA 1.49 TIRF objective. Acquisition was triggered using a Matlab script, which in turn used Micromanager for capturing images, launched WOSM macros for all other microscope functionality, and controlled and monitored the microfluidics. Pixels were 130 nm. Data were acquired at 23 °C.

### Analysis of MT shrinkage rates

Data were analysed in Matlab (Mathworks). Each MT image was aligned horizontally, using the function *imrotate* to rotate the image according to a hand-drawn line, before averaging columns of 11 pixels spanning the MT to generate an intensity profile. This was repeated for the same region of interest (ROI) for every frame in an image stack. Kymographs were produced by vertically concatenating the intensity profiles. Shrinkage rates were measured by manually tracing kymographs using the impoly function, extracting coordinates and calculating the slope. Time and distance calibration was automated by extracting the image metadata. Rates quoted in this paper assume a conversion of 125 dimer PF^−1^= 1 μM. Quantitative fluorescence analysis of MTs in a kinesin-clamp is presented in Supplementary Methods. Graph plotting and statistical tests were also carried out using Matlab.

### Analysis of MT expansion assay with microfluidics

Kymographs were generated using the Fiji/ImageJ plugin KymoResliceWide and the subsequent analysis was performed using Matlab. The expansion of the MT was measured for each time point as shown in Supplementary Figure 5. These measurements were then analysed for each ADP or kinesin interval: the mean value was taken for each ADP interval, whereas a logistic curve was fitted (using bisquare robust fitting) to each ADP-kinesin transition to estimate the kinesin-driven expansion (as shown in Fig. 5d).

### Analysis of surface-stitched MT expansion assay

Coordinates of MTs were extracted from dark-field images using the semi-automated Fiji/ImageJ plugin, JFilament^36^. The coordinates were then mapped onto the fluorescence channel and used to generate a 5-pixel-wide line scan. The fluorescence profiles of the cap and the seed were each fitted with a Gaussian error function in Matlab. The length of the GDP-bound section of the MT is then given by the distance between the point of inflection on each curve. The MT length change was then assessed by taking the mean length of the manually identified plateaus, as shown in Fig. 5c. Points deviating by greater than 5% from the median length in these intervals were discarded prior to fitting. Long MTs were selectively chosen for the analysis to improve precision, with the average length of kinesin-free GDP-MT segments being 41 μm.

## Acknowledgements

The authors thank D. R. Drummond and N. Sheppard for assistance with protein purification, and T. A. McHugh for invaluable comments on the manuscript. This research was funded by the Biotechnology and Biological Sciences Research Council (grant number BB-G530233-1) via the Systems Biology Doctoral Training Centre, University of Warwick; and the Wellcome Trust (grant number 103895/Z/14/Z).

## Author contributions

D.R.P. and R.A.C. designed experiments. N.J.B. provided mathematical insight. D.R.P. collected and analysed the data, and produced the manuscript and figures. All authors contributed towards the discussion and interpretation of results, and editing the manuscript.

## Competing financial interests

The authors declare no competing financial interests.

**Supplementary Movie 1 | A kinesin-clamp assay.** The image data (*top*) corresponds to the kymograph in Fig. 2c (*bottom*). A minus-end trail is clearly seen in the no nucleotide phase. Addition of ADP causes the MT tips to shrink. In this case, the minus-end trail is retained during shrinkage. MTs are re-stabilised upon addition of taxol and ATP, and the resulting kinesin-driven MT gliding reveals the MT polarity.

**Supplementary Movie 2 | Strong-state kinesin can lock the curvature of GDP-MTs.** For each concentration of T93N, images are sorted according to the MT orientation. The marked MTs in each row (*orange asterisks*) fall into the orientation range depicted by the protractor diagrams (*left*). MTs are straight and dynamically unstable at the beginning of the movie. Arrows (*top*) highlight the presence and direction of hydrodynamic flow, which causes MT bending. In the absence of kinesin, stopping the flow causes the MTs to re-straighten and continue to depolymerise. MT curvature is preserved at low concentrations of T93N but not at high concentrations. MTs also transiently crinkle when T93N is flowed through at high concentrations.

**Supplementary Movie 3 | Kinesin reversibly expands MTs under constant hydrodynamic flow.** The movie corresponds to the MT shown in Fig. 5b. As kinesin and ADP are alternately introduced into the flow, the MT visibly expands and contracts, most obviously seen by the downstream MT tip visibly shifting right and left. Both surface-free and surface-snagging behaviour is shown in this movie, and the MT expands to the same extent in each case (Fig. 5d).

**Supplementary Movie 4 | MTs briefly crinkle when kinesin is introduced to the flow.** An ADP-to-kinesin transition is shown, during which the MT crinkles and thereby dips in and out of the TIRF illumination. Most of the MTs are visibly free from the surface for the duration.

**Supplementary Movie 5 | Kinesin increases the lattice spacing of surface-stitched GDP-MTs.** The movie corresponds to the MT shown in Fig. 5f. Part way through the movie, 200 nM of monomeric kinesin (K340) was flowed through the channel and the MT extends and bows so as to follow a longer path length. Flushing with 1 mM ADP triggers kinesin unbinding, and the MT reverts to its original length. After washing the sample thoroughly with buffer, the process can be repeated. After the first cycle, the MT becomes tethered to the surface at a greater number of interaction sites. During the second kinesin flow-through, part of the MT briefly goes out of focus but it is recruited back into the optical plane, demonstrating that our protocol restricts motion in the *z*-axis to permit reliable quantification. Scale bar is 10 μm.

**Supplementary Animation 1 | Kymograph profile matching for measuring MT expansion.** The ‘original’ kymograph shown is the same as in Fig. 5b. The ‘transformed’ kymograph is the same image but each row has been compressed by the values shown in Fig. 5d and then the magenta imaging channel was used for alignment. This highlights the expansion that occurs when kinesin binds to the MT.

